# Comparative virulence analysis of seven diverse strains of *Orientia tsutsugamushi* reveals a multifaceted and complex interplay of virulence factors responsible for disease

**DOI:** 10.1101/2024.12.18.629091

**Authors:** Panjaporn Chaichana, Naphat Satapoomin, Chitrasak Kullapanich, Suthida Chuenklin, Aisha Mohammad, Manutsanun Inthawong, Erin E. Ball, Thomas P. Burke, Piyanate Sunyakumthorn, Jeanne Salje

## Abstract

*Orientia tsutsugamushi* is an obligate intracellular bacterium found in *Leptotrombidium* mites that causes the human disease scrub typhus. A distinguishing feature of *O. tsutsugamushi* is its extensive strain diversity, yet differences in virulence between strains are not well defined nor well understood. We sought to determine the bacterial drivers of pathogenicity by comparing murine infections using seven strains combined with epidemiological human data to rank each strain in terms of relative virulence. Murine cytokine expression data revealed that the two most virulent strains, Ikeda and Kato, induced higher levels of IL-6, IL-10, IFN-γ and MCP-1 than other strains, consistent with increased levels of these cytokines in severe human scrub typhus patients. We sought to identify the mechanistic basis of the observed differential virulence between strains by comparing their genomes, *in vitro* growth properties and cytokine/chemokine induction in host cells. We found that there was no single gene or gene group that correlated with virulence, and no clear pattern of *in vitro* growth rate that predicted disease. However, microscopy-based analysis of the intracellular infection cycle revealed that the only fully avirulent strain in our study, TA686, differed from all the virulent strains in its subcellular localisation and expression of its surface protein ScaC. We conclude that drivers of pathogenicity in *Orientia tsutsugamushi* are distributed throughout the genome, likely in the large and varying arsenal of effector proteins encoded by different strains, and that these interact in complex ways to induce differing immune responses and thus differing disease outcomes in mammalian hosts.

**Author Summary:** Scrub typhus is a vector-borne human disease caused by the bacterium *Orientia tsutsugmushi* and spread by mites. There are numerous different strains of this bacterium with some causing more severe disease in humans than others, and some that do not cause any illness at all. The factors driving these differences are not yet understood, and gaining insight into them could aid in vaccine development and help predict the severity of disease caused by new isolates. To better determine the mechanistic basis of pathogenicity in scrub typhus, we carried out experiments in which we compared seven diverse strains for virulence in animals. We measured their ability to cause disease in mice, so that we could reliably classify them as virulent or avirulent in this model. We then analysed various genomic and biological aspects to identify disease markers in both mice and humans. We report here that there is no single factor that predicts whether a strain will be pathogenic or not, but that disease in scrub typhus is a complex process resulting from the activity of multiple bacterial genes working together to drive different immune responses in the host, resulting in either clearance of the bacteria from the host, or escalating disease. Future work exploring the relationship of bacterial effector proteins will help to disentangle this complex relationship in mechanistic detail.

## Introduction

Scrub typhus is a severe vector-borne human disease caused by the obligate intracellular bacterium *Orientia tsutsugamushi* (Ot, Order Rickettsiales, Family Rickettsiaceae)^1–3^. Symptoms include headache, fever, and rash, and if not treated promptly and with effective antibiotics this can escalate to multiple organ failure and death. Scrub typhus is a leading cause of severe febrile illness in many parts of Asia, particularly in rural regions, while related species causing scrub typhus-like disease have recently been described in other parts of the world, namely *Candidatus* Orientia chiloensis in Chile^4–6^ and *Candidatus* Orientia chuto in the Middle East^7^.

While Ot can cause disease in humans, its primary reservoir and vector is the *Leptotrombidium* mite. Ot is found in the salivary glands and ovaries of these arthropods. It can be transovarially transmitted, allowing it to maintain its presence in mites without the need for an animal reservoir. Mites feed on rodents and other small mammals during the larval stage of its lifecycle, and Ot can be detected in these animals in the wild^8–10^. Whilst Ot can be transmitted from mammals to naïve mites, the efficiency is very low and novel transovarial transmission is not easily established^11^, leading to the conclusion that mites serve as the primary reservoir. Therefore, although Ot is a human pathogen, it has primarily evolved as a reproductive parasite of mites. Most of the strains of Ot that are studied were originally isolated from human patients or small animals and it is not known whether there is a separate large population of strains found in mites that are incapable of causing disease in humans. In the closely related genus Rickettsia, which includes various tick-borne human pathogens, recent metagenomic analyses have uncovered many novel Rickettsia species in ticks^12,13^. This suggests that most Rickettsia species have primarily evolved as endosymbionts of ticks, with only a small subset capable of causing disease in humans. It is unknown whether, by analogy, there are numerous yet undiscovered Ot strains that are only found in mites, but two lines of evidence point to this. First, target capture enrichment sequencing analysis showed diverse Ot strains in mites in Southeast Asia that are absent in human populations^14^. Second, an Ot-like species was recently reported from mites in South Carolina, USA, where no cases of scrub typus like illness have been described^15^. Thus, it is possible that large numbers of uncharacterized, nonpathogenic strains of Ot exist, reinforcing the notion that pathogenesis is a variable trait that is not integral to the biology of Ot.

A key outstanding question in the field is: what bacterial factors drive disease outcomes in scrub typhus? Since genetic tools for Ot are currently unavailable, we compared the virulence of seven different isolates to identify bacterial factors influencing disease outcomes. The strains used in our study are Karp, Kato, Gilliam, UT76, Ikeda, TA686 and TA763^16^. We additionally used UT176 in some experiments to enable comparisons to our previous work that used this strain^17^. Both TA686 and TA763 were originally isolated from animals^18^, whilst all the other isolates originate from human patients. Karp, Kato and Gilliam are the three main serotypes found in Southeast Asia^18,19^, UT76 and UT176 were isolated from patients in Northern Thailand^20^ and Ikeda is a strain prevalent in Japan^21^.

The determinants of virulence for a particular strain must be encoded within the bacterial genome. Ot has an unusual genome of 2.1-2.8 Mbp, in which around 50% of the genome is comprised of multiple copies of an amplified and highly degraded mobile genetic element, the integrative and conjugative element named the rickettsial amplified genetic element (RAGE)^16,22,23^. All strains used in the current study have complete genome sequences available and we recently carried out a detailed analysis of these genomes^24^. We showed that the sequences in between the RAGEs, called inter-RAGEs (IRs) are highly conserved between Ot strains, although the ordering of the groups of IR genes along the genome varies substantially. All genomes encode 76-93 regions of RAGEs, although these are mostly heavily degraded with some as short as three genes in length. Whilst the classes of genes encoded in the RAGEs are consistent between genomes, there is substantial variation in the numbers of different genes. In particular, Ot encodes large numbers of Ankyrin repeat containing proteins (Anks) and tetratricopeptide repeat containing proteins (TPRs), both of which are predicted effector proteins. There is enormous diversity in the number and sequence of these proteins between strains. Whilst the biological function of certain Anks has been determined^25–30^, and the subcellular localisation of all the different Anks in strain Ikeda have been shown^25^, there remain the vast majority for which no function has been determined and this cannot be predicted from sequence alone. In summary, Ot genomes are complex and some virulence genes have been described, however, the genetic determinants of pathogenicity in the genomes of Ot strains remains unknown.

A comparison of strain-specific differences in the intracellular lifecycle of Ot have not been reported. Ot enters host cells using a combination of clathrin-mediated endocytosis and macropinocytosis^31,32^, whereupon it escapes the endolysosomal pathways and is located directly in the host cytoplasm. It traffics to the perinuclear region using dynein-dependent motility along microtubules^33^, driven by an interaction between the bacterial surface autotransporter protein ScaC and the dynein adaptor proteins BicD1/BicD2^34^. Ot undergoes bacterial replication in the perinuclear region over a period of 3-7 days. It then exits infected cells by relocating to the host cell surface and budding off, encased in host plasma membrane^35^. This mode of exit has not been described in other bacteria, and it is associated with a developmental transition to a distinct extracellular form of Ot^31^. Whilst aspects of this intracellular infection cycle were discovered in different strains, reflecting the main strain used by different laboratories, a comparison of the similarities and differences in the infection cycle between different strains of different virulence has not been reported.

This study investigates the mechanistic basis of pathogenicity caused by Ot. We carried out a side-by-side comparison of the relative virulence of seven diverse Ot strains in a murine model of disease and combined this with analyses of the *in vitro* growth properties of the same strains in multiple cell types. We also measured cytokines elicited by each strain *in vitro,* performed genome comparisons of virulence genes, and characterized the intracellular lifecycles of these strains by fluorescence microscopy. Together, these studies suggest that virulence is not encoded by a single gene, or group of genes, but is distributed across the genome of Ot and that disease severity results from a complex interplay between Ot factors and the host immune response to this infection.

## Results

### A comparative analysis of seven Ot strains in a C57Bl/6J murine infection model reveals a hierarchy of relative virulence

It is unknown why some Ot strains are more pathogenic than others. To address this, it was important to first determine whether the strains differ in virulence and therefore we conducted a series of murine infection experiments. Previous comparative studies using murine infection models have examined only two or three strains, making it difficult to rank a larger number by virulence^17,36–38^. Of the strains used in our study, studies have shown that Karp, Kato, and UT76 are virulent, while Gilliam’s virulence varies with the mouse strain, classifying it as intermediate.^17,36,38^.

We performed two experiments, one with a high infection dose of 10^7^ per mouse and a shorter duration of 8 days, and a second with a lower infection dose of 1.25 x 10^6^ per mouse and a longer duration of 12 days (**Fig. 1A, B**). Both experiments used 6-9 week old C57Bl/6 mice and intravenous inoculation. Ikeda and Kato caused significant weight loss and lethality in the high dose experiment, with UT76 also causing low but significant weight loss over the course of the experiment (**Fig. 1A, B**). In the low dose experiment, only Kato caused lethality (**Supplementary Fig. 1A**) but Kato, Ikeda and UT76 all caused significant weight loss after 7 days (**Fig. 1C**). By comparison, lethality and significant weight loss was not observed in any other strain in either experiment (**Fig. 1B, C**). The bacterial levels in the blood at the time of termination were correlated with weight loss and lethality in the high-dose experiment, with UT76, Ikeda, and Kato exhibiting the highest levels of bacteremia. (**Fig. 1D**). In the low-dose experiment, although UT76, Ikeda, and Kato all exhibited the highest levels of bacteremia, UT76 levels were significantly higher than those of the other two strains, indicating that the animals were less able to clear this particular strain under these experimental conditions. (**Fig. 1E)**.

**Figure 1.**
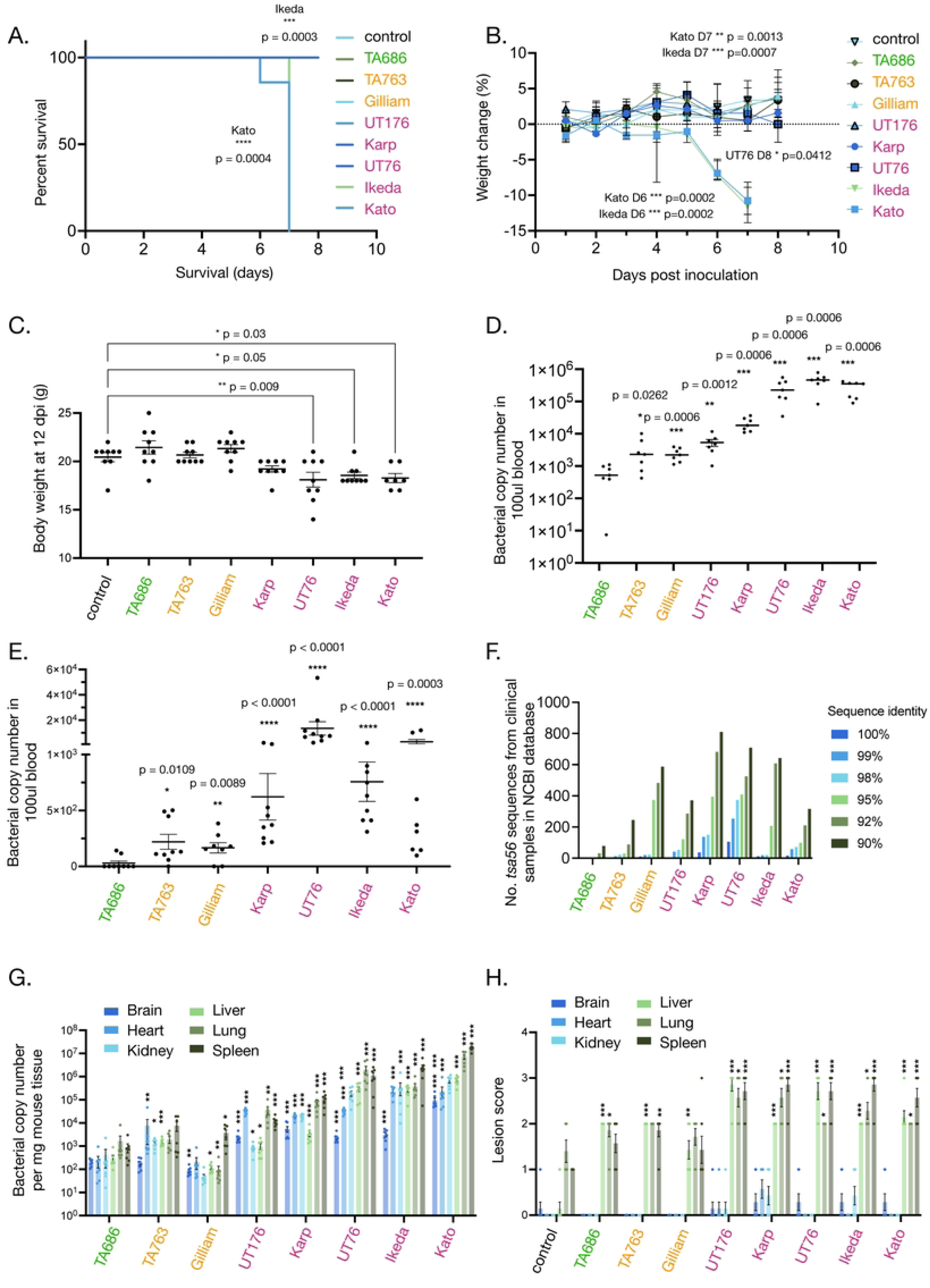
Ot strains can be classified into avirulent, intermediate virulent, and virulent groups. A-B. Results from high dose experiment in which C57Bl/6J were infected with 1x10^7^ bacteria per animal and monitored for 8 days. A. Kaplan-Meier survival curves of mice infected with different bacterial strains showing that strains Kato and Ikeda cause lethal infection in this animal model. Statistical significance was determined using log-rank Mantel-Cox analysis. B. Graph showing percentage weight change over time, whereby only strains Kato, Ikeda and UT76 cause significant weight change during the course of infection. Changes in body weight of Ot inoculated groups were compared to mock control on specific days by One-way ANOVA followed by Dunn’s multiple comparisons test. C. Results from low dose experiment in which C57Bl/6J were infected with 1.25x10^6^ bacteria per animal and monitored for 12 days. Graph showing body weight at the point of termination whereby only strains UT76, Ikeda and Kato cause significant weight loss. Mean and SEM are shown. Body weight of Ot inoculated groups were compared to mock control by One-way ANOVA followed by Dunnett’s multiple comparisons test. D-E. Bacteremia in high dose (D) and low dose (E) experiment. Graph shows bacterial load in mouse blood at point of termination as measured by qPCR using primers against the conserved single copy bacterial gene *tsa47.* Mean and SEM are shown. Statistical significance was determined using Mann Whitney test in which each strain was compared with TA686. F. Numbers of DNA sequences of the Ot-specific gene *tsa56* available on the NCBI database on 1 July 2024 with 90%, 92%, 95%, 98% and 100% sequence identity to the *tsa56* sequence from different Ot strains used in our study. Sequences in the NCBI database are almost entirely derived from clinical samples, so this analysis reflects the global levels of strains related to the strains in our study present in reported clinical studies. G. Bacterial load in different mouse tissues at the point of termination as measured by qPCR using primers against the conserved single copy bacterial gene *tsa47.* Mean and SEM are shown. Statistical significance was determined using Mann Whitney test in which each strain was compared with TA686. * p ≤ 0.05, ** p≤ 0.01, *** p ≤0.0001. H. Tissue damage caused by different strains in different tissue types as assessed by lesion score analysis of H&E-stained sections. Mean and SEM are shown. Statistical significance was determined using Mann Whitney test in which each strain was compared with uninfected control. * p ≤ 0.05, ** p≤ 0.01, *** p ≤0.0001. Representative images are given in Supplementary Figure 1.

We sought to compare these data with previous studies to better categorise the seven strains. A study using silvered leaf monkeys compared the relative virulence of Karp, Kato, Gilliam, TA678 and TA686^39^. It was shown that Karp, Kato and Gilliam caused illness in these animals, with some lethality observed in both Karp and Gilliam. By contrast, TA678 induced only limited clinical symptoms, and TA686 no clinical symptoms at all as measured by fever, lymphadenopathy, eschar formation or leucocytosis. Comparing disease outcomes in human patients infected with different strains is challenging due to factors like underlying health, initial bacterial load, and treatments received. However in a study in Hainan Island, Southern China, higher bacterial loads were observed in patients infected with Karp compared to Gilliam, while patients infected with TA763 had undetectable levels^40^. Similarly a study in South Korea showed more severe clinical features in patients infected with a South Korean isolate Boryong compared with Karp^41^. We carried out an analysis of the phylogeny of available *tsa56* gene sequences in the NCBI database to establish the distribution of strains reported from humans (**Fig. 1F**). This gene is typically used to classify Ot strains, and most of the sequences deposited in this database are derived from human clinical studies. Therefore, this analysis gives some indication of the abundance of different strains causing disease in humans. Whilst the distribution of strains is biased by the geographical location of research activity, it is notable that the numbers of strains with >90% identity to Gilliam was close to that of Ikeda, UT76 and Karp. Out of 4,710 sequences analyse there were none identified with ≥98% identity to TA686, suggesting this strain is not found in humans.

Based on these data, we classify Kato as the most virulent strain, followed by Ikeda and then UT76. Since Gilliam and TA763 did not cause weight loss or lethality, but are found in human patients, we classify these as strains with intermediate virulence, as has been described before for Gilliam. Finally, the fact that TA686 had no significant weight loss or mortality, combined with its demonstrated lack of virulence in a non-human primate study and its absence from databases of human patient isolates (**Fig. 1F**), suggests that this strain is avirulent.

### Different Ot strains exhibit different tissue tropisms

The tissue tropisms of the different Ot strains remained unclear. We compared the tissue tropism of eight Ot strains by carrying out qPCR analysis of the brain, heart, kidney, lung, liver and spleen of all the animals in the high dose experiment (**Fig. 1G**) and combined this with an H&E staining analysis of the inflammation and tissue damage observed in the same tissues (**Fig. 1H, Supplementary Fig. 1B-E**). Both bacterial levels and tissue damage were generally lower in the avirulent and intermediate virulent strains compared with the virulent strains. Within the organs the brain showed the lowest bacterial loads with mean 87 – 9.5x10^4^ Ot copies per mg, and lung and spleen the highest with mean 94 - 7x10^6^ and mean 880 – 1.9x10^7^ Ot copies per mg respectively, whilst tissue damage as measured by lesion scoring was low in brain, heart and kidney tissue (mean lesion scores 0-0.3, 0-0.6 and 0-0.4) but high in liver, lung and spleen (mean lesion scores 1.4-2.8, 1.7-2.6 and 1.4-2.9). There were some notable strain-specific outliers. UT176 was particularly abundant in the heart compared with other tissues and other strains, whilst Kato was comparatively abundant in the brain. In terms of inflammation, Karp exhibited the highest inflammation and tissue damage in the brain, heart and kidneys followed by UT176. These data indicate that strain-specific differences in virulence do not follow a linear scale; instead, each strain interacts uniquely with the host, resulting in distinct outcomes.

### Ot strains elicited distinct immune responses in a murine infection model

The main driver of pathogenesis in human disease is an overactivation of the inflammatory response leading to disseminated vascular coagulation and tissue damage^42,43^. We previously compared two strains, Karp and UT176 (distinct from UT76 used in the current study) and found that Karp caused greater disease severity in a mouse model of infection, and induced more proinflammatory cytokines including IL33 in a dual RNAseq analysis of Ot-infected HUVEC cells^17^. A different study compared the virulence of Karp and Gilliam and two murine species, C57Bl6 and outbred CD1 and found that both caused disease in CD1 but only Karp caused disease in C57Bl/6^38^. In the C57Bl/6 mouse strain, Karp caused a stronger immune activation, including infiltration of CD4+ and CD8+ T cells and increased serum cytokine/chemokine levels compared with Gilliam.

We sought to explore how the multiple different Ot strains caused inflammation *in vivo*. We used Luminex profiling of serum taken at 6 days post infection in the high dose cohort to analyse the relative levels of 9 cytokines and chemokines: IL-1α, IL-1ß, IL-6, IL-10, IL-12(p70), IL-17, CCL2/MCP-1, GM-CSF and IFN-γ (**Fig. 2**). MCP-1, IL-6 and IFN-γ exhibited a pattern that strongly correlated with virulence, whereby the avirulent and intermediate virulence strains TA686, TA763 and Gilliam induced low levels, while the virulent strains UT76 and Karp induced high levels, and high virulence strains Kato and Ikeda induced very high levels of cytokine. IL-10 showed the same pattern although no cytokine was detected in TA763. The virulent strain UT176 showed no induction of IL-17, IL12(p70), IL-10, IL-6, IL-1ß, IL-1a and IFN-γ suggesting a generally reduced inflammatory response relative to other strains. GM-CSF was upregulated relative to the control to a similar extent in all strains, as was IL-1a with the exception of UT176 that did not induce any detectable cytokine. IL-17, IL-12(p70) and IL-1ß were differentially upregulated in different strains without any patterns that correlate with virulence. Together, these data show that different strains lead to different signatures of inflammatory responses *in vivo*, but that relative induction of MCP-1, IL-6, IL-10 and IFN-γ correlates with degree of virulence in a murine infection model. Previous analysis of severe scrub typhus in human patients have shown that these same cytokines and chemokines are associated with severe disease in humans.

**Figure 2.**
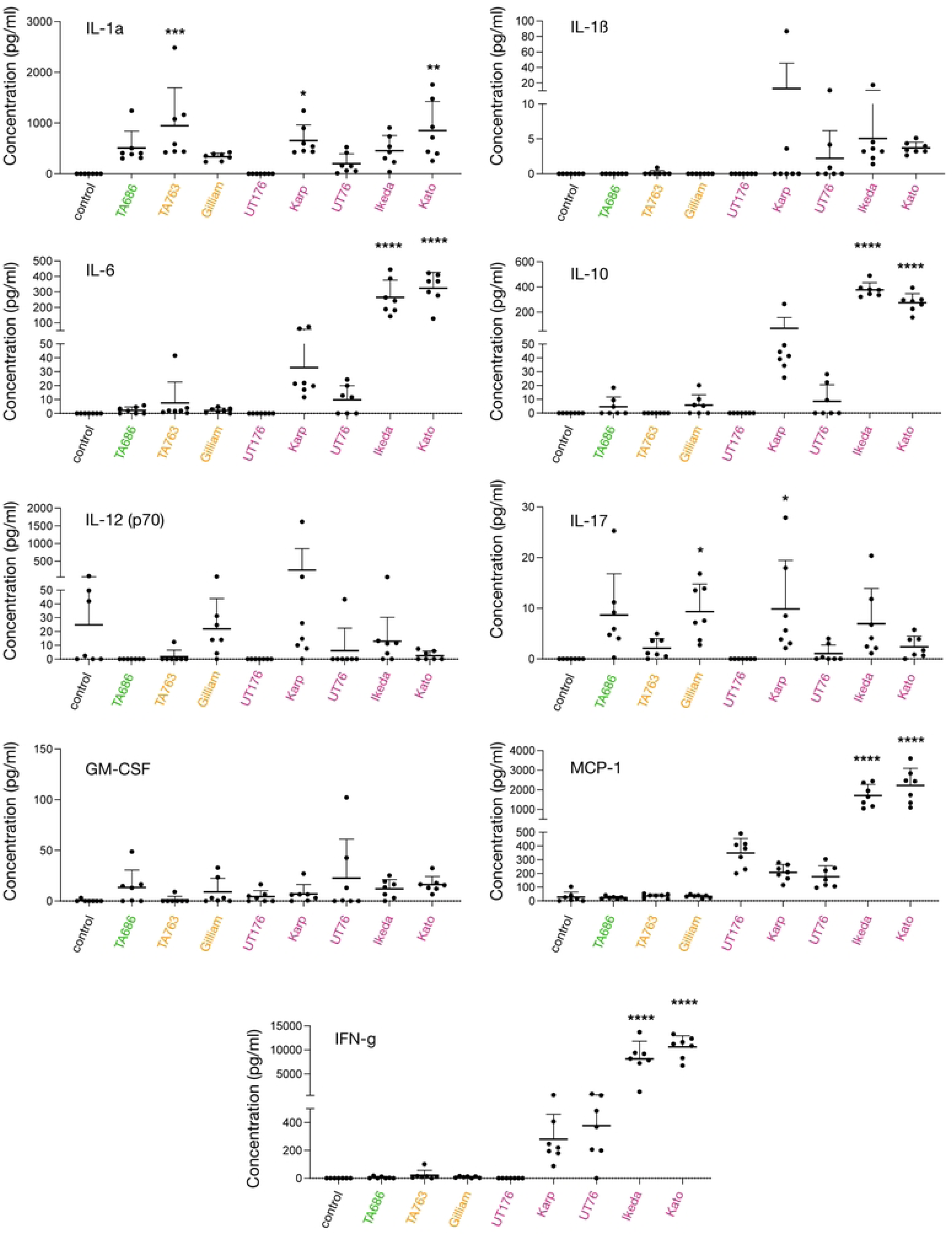
Cytokine profiling reveals a positive association of IL-6, IL-10, IFN-γ and MCP-1 with virulence in a murine infection model. Graphs show the relative levels of nine chemokines and cytokines in mouse blood collected from the tail vein at six days post infection in the high dose experiment described in Figure 1, as measured by Luminex analysis. Statistical significance was determined using one way ANOVA test followed by Tukey’s multiple comparisons test in which all groups were compared with mock infected control. * p<0.05, *** p<0.005, **** p< 0.0001.

### Comparative genomics reveals complexity of potential virulence genes in Ot

We sought to identify genetic determinants of pathogenicity in the genomes of Ot strains. First, we compared the RAGE regions of the Ot genome, that are composed of a highly proliferated integrative and conjugative element and make up around 50% of the Ot genome. We were not able to identify any patterns in the number or completeness of RAGEs that correlate with virulence (**Fig. 3**). We then examined the distribution of predicted effector proteins of the Ankyrin repeat (Ank) protein families, that are abundant within Ot RAGEs. We examined a published analysis of the 54-75 Ank genes present across Ot strains and could not identify any patterns of distribution that correlated with virulence. However, there is extensive diversity in the arsenal of Anks encoded by Ot genomes, and function cannot be predicted by sequence. It is therefore highly likely that the unique complement of Ank proteins encoded by each Ot strain results in specific manipulation of host cells, leading to different outcomes in terms of bacterial clearance or disease progression.

**Figure 3.**
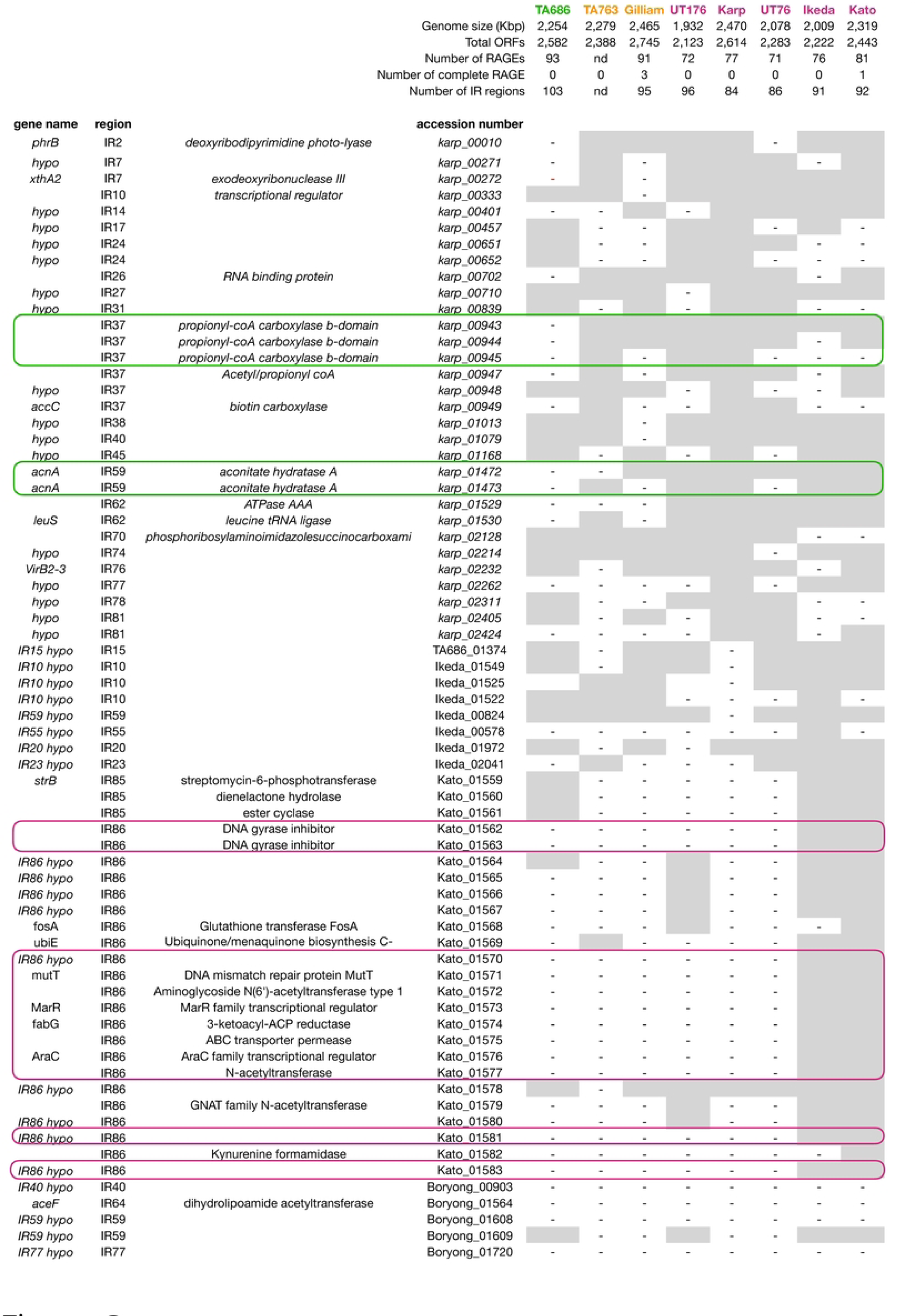
Identification of genes whose presence/absence correlates with virulence. The presence or absence of any inter-RAGE genes not uniformly present across all eight genomes included in this analysis are shown. Green lines highlight genes uniquely absent in TA686 whilst magenta lines highlight those genes uniquely present in Ikeda and Kato. The inter-RAGE region in which the gene is located is shown in column 2. Additional information about each genome is given at the top of the table. ORF = open reading frame, RAGE = Rickettsial Amplified Genetic Element, IR = Inter-RAGE region.

We then compared the non-RAGE regions of the different Ot strains to identify genes uniquely present or absent in pathogenic strains. The non-RAGE genes are encoded in small groups called inter-RAGE blocks (IRs) that are non-syntenous between strains due to the presence of very high numbers of RAGEs. We observed that the avirulent strain TA686 contains the largest number of IRs, due to increased fragmentation of IRs compared to other genomes. However, the total number of virulent and avirulent genomes in our analysis is not sufficient to determine whether this is a virulence-related correlation. We analysed the identity of IR genes and identified 657 core genes present in all Ot genomes, and an additional 74 IR genes missing in one or more genomes. We analysed all 74 to seek patterns that correlate with virulence (**Fig. 3**). We identified 2 genes uniquely absent in the avirulent strain TA686. These were propionyl-coA carboxylase b-domain gene and aconitate hydratase A. These genes are not known to be associated with pathogenicity in other organisms, although it is possible that their absence in TA686 is associated with a decreased survival *in vivo*. A lack of genetic tools makes it difficult to further test the relevance of these genes in virulence at the current time. We did not identify any genes that were present in all virulent strains compared with those of intermediate or no virulence. However, there were 19 genes uniquely found in Kato and Ikeda, the two most virulent strains. All but one are located in a single IR block: IR86, present only in these two strains. It is possible that the presence of this additional block of genes leads to increased bacterial growth and/or pathogenesis *in vivo.* However, it is important to note that Kato and Ikeda are more closely related to each other than to other strains in our study and therefore it is possible that this difference reflects genetic relatedness and has no impact on virulence. A larger number of genomes as well the availability of genetic tools will enable further studies on the relevance of these genes.

### Ot strains grow at similar rates in macrophages and other cell types in vitro

In the case of the sister genus, *Rickettsia,* an inability to grow in macrophage cells is associated with a lack of pathogenicity^44,45^, and we therefore hypothesized that *in vitro* growth rates of Ot strains would vary in different cell types in vitro. We measured the growth rate from 4-7 days in a mouse fibroblast cell line L929, a mouse macrophage cell line Raw264.7, mouse bone marrow derived macrophages (BMDMs), a dog macrophage cell line DH82, a human macrophage cell lines THP1, and human derived primary macrophages (**Fig. 4A)**. The growth rates were highly similar between strains, with no significant differences that correlated with virulence. These data suggest that differential virulence in Ot is not determined by intrinsic differential bacterial growth rates in certain cell types, unlike related pathogens.

**Figure 4.**
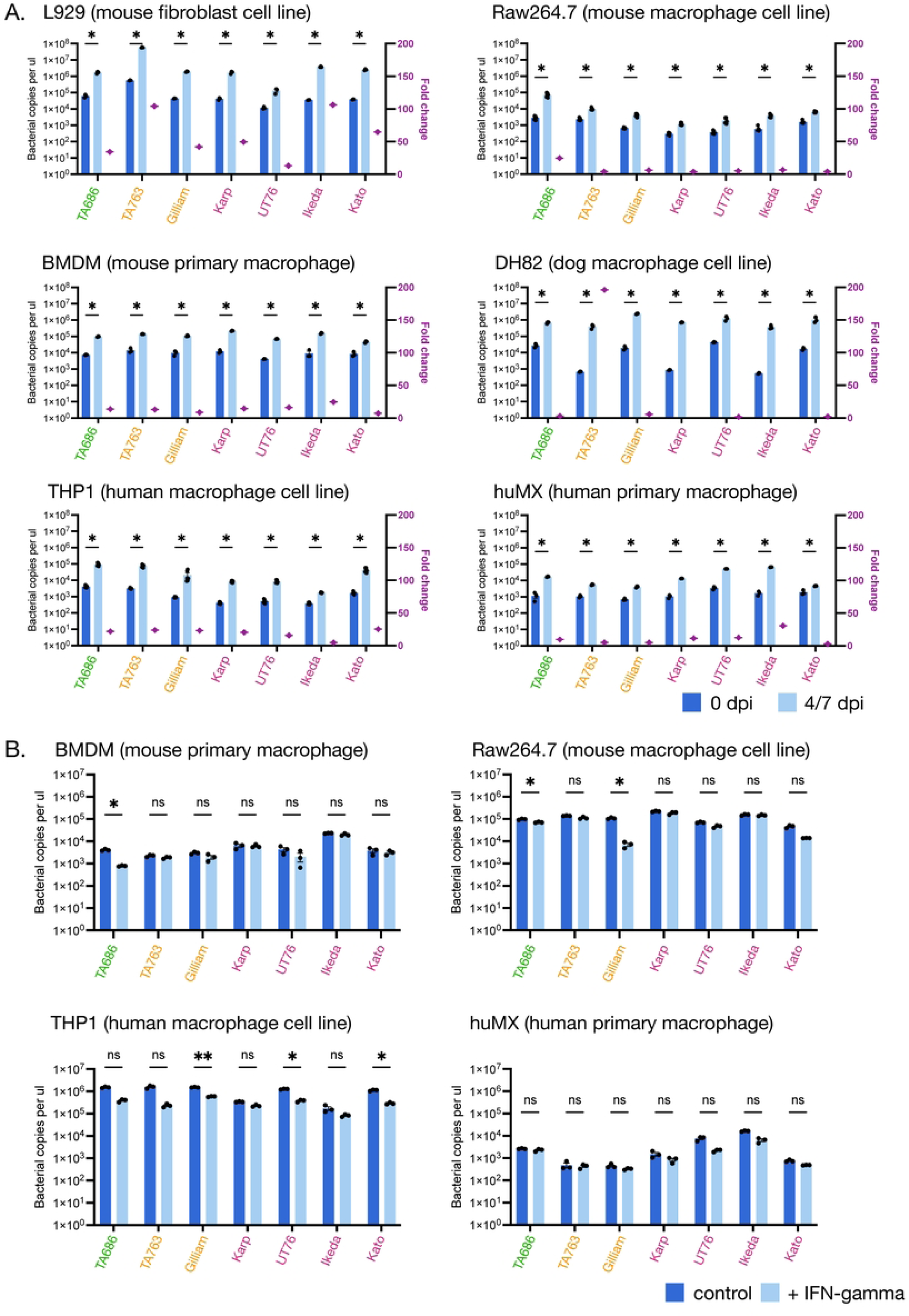
The *in vitro* growth rates of different Ot strains are similar to each other. A. Growth of seven strains in L929, Raw264.7, BMDM, DH82, THP1 and huMX cells. Bacteria were grown in different cell lines for 7 days (DH82) or 4 days (other cell lines). Bacterial levels were measured by qPCR using primers against the Ot-specific gene *tsa47.* Data are shown as mean ± SEM (n = 6); statistical significance of growth increase for each strain was calculated using t-test where *, p<0.05; **, p<0.01, ***, p<0.005. Additional statistical tests comparing the significance of the fold increase between all strains in one cell line were determined using a one-way ANOVA followed by Tukey’s multiple comparisons, and these results of these tests are given in Supplementary data X. B. Growth of seven strains in BMDM, Raw264.7, THP1 and huMX cells in the presence and absence of IFN-γ.Bacteria were grown in different cell lines for 4 days. Bacterial levels were measured by qPCR using primers against the Ot-specific gene *tsa47* and the graph shows the bacterial level after 4 days growth in the presence or absence of 100U/ml of recombinant mouse or human IFN-γ. Data are shown as mean ± SEM (n = 3). The statistical significance of the difference between growth in the presence or absence of IFN-γ across strains within one cell line was determined using two-way ANOVA followed by Bonferroni’s multiple comparisons, where *, p<0.05; **, p<0.01.

### Differential growth in interferon-activated macrophages does not explain differential virulence in Ot

IFN-γ is an anti-microbial cytokine that is upregulated in scrub typhus patients and was recently shown to be critical for controlling Ot *in vivo*^46^, as well as *Rickettsia parkeri* in macrophages *in vitro*^47^ and *in vivo*^48^. We therefore hypothesized that it may differentially restrict Ot strains *in vitro* based on their virulence *in vivo*. We measured intracellular bacterial abundance in the presence or absence of IFN-γ in four macrophage cell lines and primary cell types (**Fig. 4B)**. We found that stimulation with IFN-γ caused either no change or a slight decrease in bacterial growth in BMDMs and RAW264.7s. However, the observed differences did not correlate with measured differences in virulence *in vivo.* We therefore concluded that differential ability of Ot strains to replicate in IFN-γ stimulated macrophages is not the primary driver of differential virulence *in vivo*.

### Cellular activation by IFN-γ leads to differences in the relative chemokine and cytokine induction by different strains of Ot, but these do not correlate with virulence

Our *in vivo* findings showed that the more virulent strains induced higher levels of IL-6, IL-10, IFN-γ and MCP-1 (**Fig. 2**). This motivated us to determine whether we could determine any differences in chemokine and cytokine production *in vitro* that correlate with virulence. Therefore, we measured the production of cytokines and chemokines in mouse (**Fig. 5A)** and human (**Fig. 5B)** primary macrophages, both in the presence and absence of IFN-γ. We used RT-qPCR to determine the relative transcript levels of IL-6, TNF-alpha, IL-1ß, IL-33, CCL2/MCP-1, CCL4/MIP-1b, CCL5/RANTES, CXCL9 and CXCL10 in both uninfected cells and cells infected with seven different Ot strains, in the presence or absence of IFN-γ. We found that infection with all strains of Ot upregulated the expression of IL-6, TNF-alpha, IL-1ß, and CXCL-10 (human macrophages only) both in the presence and absence of IFN-γ. We observed that for TNF-alpha, IL-33, CCL5/RANTES in mouse macrophages and IL-6, TNF-alpha, CCL2/MCP-1, CCL4/MIP-1b and CXCL9 in human macrophages, all Ot strains supressed the relative increase in expression upon IFN-γ expression. Exceptions from these trends included a high induction of mouse IL-6, IL-1B, IL-33 and human TNF-α and CCL5/RANTES by Ikeda, and high induction of mouse CCL2/MCP-1 by TA686. Taken together, the analysis revealed a heterogenous distribution of relative induction of cytokines and chemokines by different strains, differential effect of IFN-γ stimulation on different strains, and differential results in different cell lines and host species.

**Figure 5.**
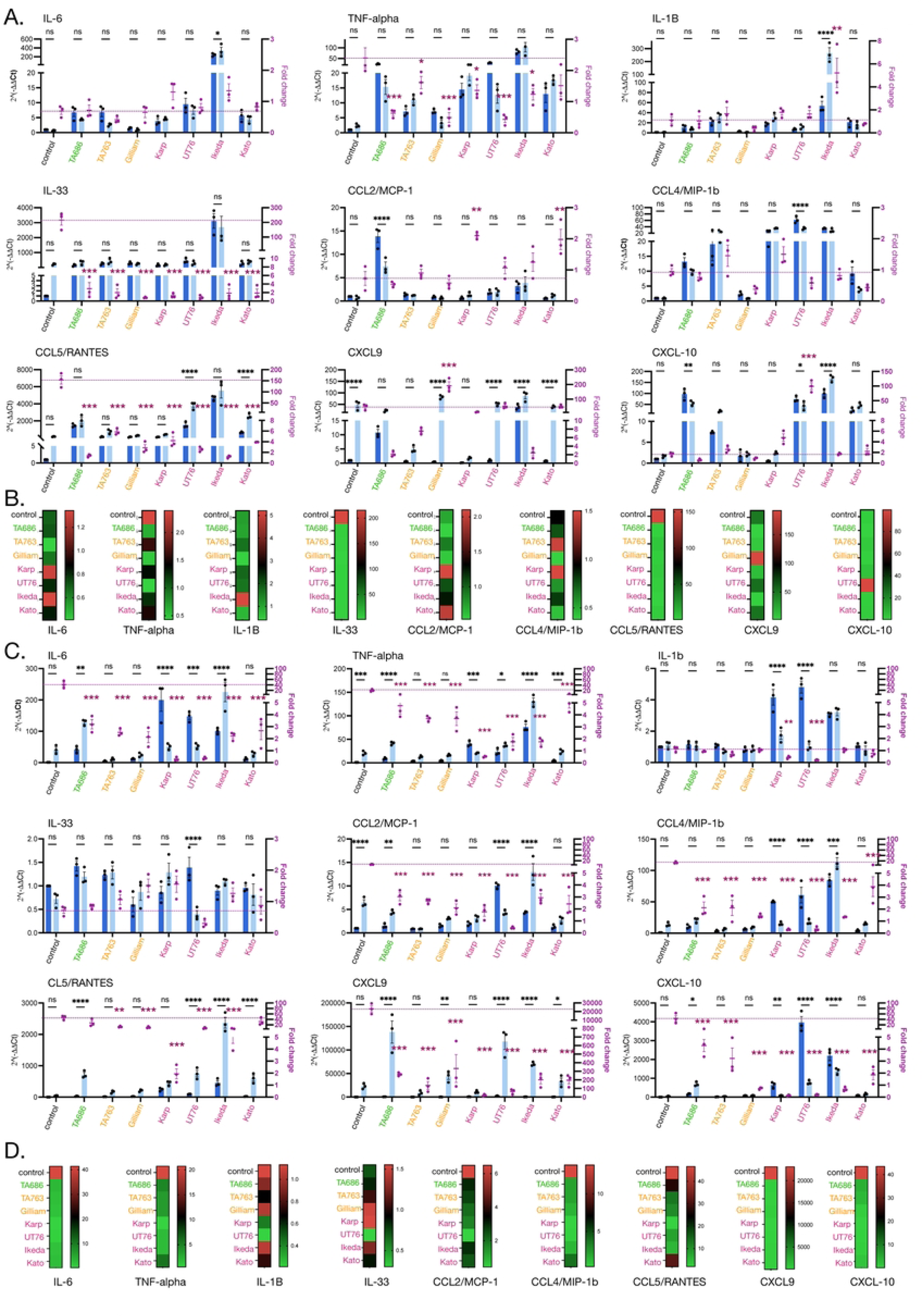
Different Ot strains induce different patterns of chemokine and cytokine induction in cultured macrophages. Graphs (A) and heatmaps (B) showing relative induction of nine chemokines and cytokines in response to infection with seven strains of Ot in murine bone marrow derived macrophages. Bacteria were infected in murine bone marrow derived macrophages for 4 days in the presence or absence of IFN-γ. The RNA was extracted and subjected to RT-qPCR using primers specific to different cytokines/chemokines. A. 2^(-1Δ1ΔCt)^ (left Y-axis) and fold changes between the absence and presence of IFN-γ (right Y-axis). The statistical significance of the difference between cytokine/chemokines genes in the presence or absence of IFN-γ across strains was determined using two-way ANOVA followed by Bonferroni’s multiple comparisons, where *, p<0.05; **, p<0.01***, p<0.001. Additional statistical tests comparing the significance of the fold change of cytokine gene expression in IFN-γ-treated infected cells compared with uninfected control was determined using two-way ANOVA followed by Tukey’s multiple comparisons. Graphs (C) and heat maps (D) showing relative induction of nine chemokines and cytokines in response to infection with seven strains of Ot in human primary macrophages. Bacteria were infected in human primary macrophages for 4 days in the presence or absence of IFN-γ. The RNA was extracted and subjected to RT-qPCR using primers specific to different cytokines/chemokines. A. 2^(-1Δ1ΔCt)^ (left Y-axis) and fold changes between the absence and presence of IFN-γ (right Y-axis). B. Heatmap. The statistical significance of the difference between cytokine/chemokines genes in the presence or absence of IFN-γ across strains was determined using two-way ANOVA followed by Bonferroni’s multiple comparisons, where *, p<0.05; **, p<0.01, ***, p<0.001. Additional statistical tests comparing the significance of the fold change of cytokine gene expression in IFN-γ-treated infected cells compared with uninfected control was determined using two-way ANOVA followed by Tukey’s multiple comparisons.

### The avirulent strain TA686 differs from other strains in subcellular localisation and expression of surface autotransporter protein ScaC

It remained unclear if the intracellular infection cycles of Ot differed between strains. We compared the number and localization of Ot in cultured mouse fibroblast cells (L929) at four days post infection, at which point bacteria are undergoing active replication, and observed differences in subcellular localisation in TA686 compared with other strains (**Fig. 6A**). While other bacteria were primarily located in a tight clump in the perinuclear region, or a mixture of tight perinuclear clump and diffuse cytoplasmic location, TA686 has a distribution that is primarily in the cytoplasm with 54% of infected cells exhibiting bacteria dispersed throughout the cytoplasm, compared with 4-21% of cells in other strains (**Fig. 6B**). We repeated the comparison between Karp and TA686 in three other cell lines, human umbilical vein endothelial primary cells HUVEC, human retinal pigment epithelial cells Rpe1 and HeLa cells, and observed the same trend (**Supplementary Fig. 2).**

**Figure 6.**
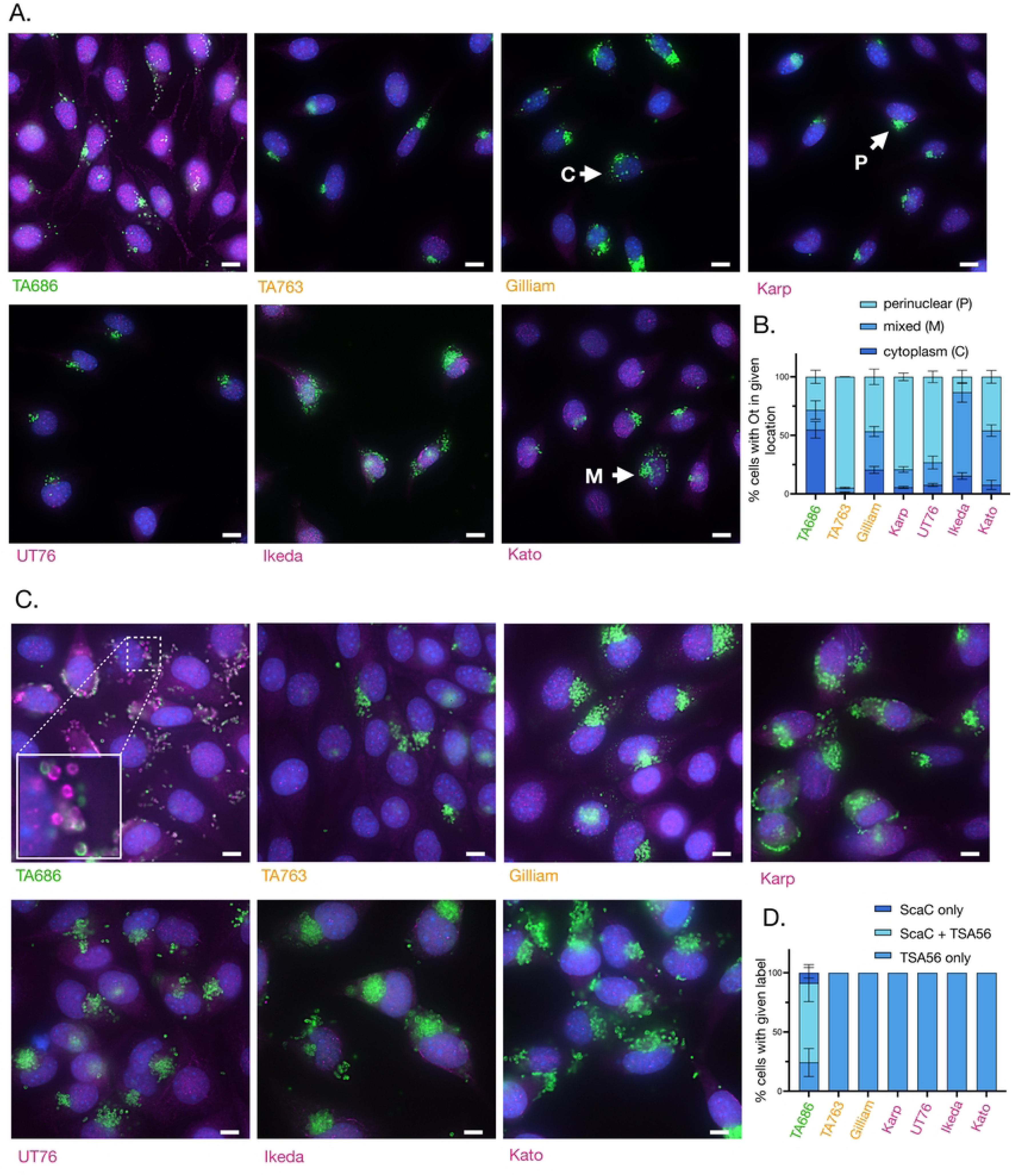
TA686 differs from other strains in its subcellular localisation and ScaC expression. Immunofluorescence microscopy images of seven strains of Ot infected with the same MOI and grown for 4 (A. and B.) or 7 (C. and D.) days in cultured L929 cells. Infected cells were fixed and labelled with an antibody against the abundant surface protein TSA56 (green) and the surface autotransporter protein ScaC (magenta). Host cell nuclei were labelled with Hoechst (blue). Representative images are shown in A. and C. whilst quantification is given in B. and D. Quantification was performed by manually scoring at least 50 infected cells from three independent replicates. In B. infected cells were scored as having bacteria in a perinuclear cluster, dispersed throughout the cytoplasm, or in both a perinuclear cluster and dispersed through the cytoplasm. In D. infected cells were scored as having bacteria inside them that were labelled with TSA56, ScaC, or both.

We then hypothesized that the surface protein ScaC might be expressed differently across these strains and examined its expression patterns. We chose this protein as a marker for the intracellular infection cycle because previous research indicated it was found on only a small fraction of bacteria, primarily located outside host cells^31^. Thus, this differentially expressed protein could serve as a useful indicator for observing phenotypic differences between strains. We observed that there was very low ScaC expression on intracellular bacteria in six Ot strains grown in L929 cells, consistent with our extensive previous studies on this protein. By contrast, TA686 expressed high levels of ScaC in a large proportion of its bacterial cells at seven days post infection (**Fig. 6C, 6D**). Together, these data show that the intracellular infection cycle of avirulent strain TA686 differs from all other Ot strains included in our study, though the relationship between these observations and virulence, as well as their generalizability to other avirulent strains, are unknown.

## Discussion

Scrub typhus is a severe human disease that is caused by a multitude of diverse Ot strains. However, the drivers of human disease in this bacterium, and the reason why some strains are more virulent than others remains unknown. The major conclusion of this study is that each strain of Ot has a unique fingerprint of interactions with its host that results in different pathogenic outcomes in a manner that cannot currently be predicted from the bacterial genome sequence. A key unresolved question is whether analysing a larger number of Ot strains would reveal broad patterns, or if the diversity within Ot populations necessitates individual study for each new strain. We observed no obvious patterns in *in vitro* growth rate or inflammatory response that correlate with virulence, suggesting a complex interplay between multiple bacterial and host factors. However, we identified four chemokines and cytokines that correlate with virulence in a murine *in vivo* infection model: MCP-1, IL-6, IL-10 and IFN-γ. This finding is consistent with human disease studies and confirms these molecules as markers of severity in scrub typhus.

It has previously been shown that the high levels of IFN-γ produced during Ot infection is essential for controlling disease, with a lack of IFN-γ in a murine model leading to lethal infection in an otherwise non-lethal model^46^. We examined the effect of IFN-γ stimulation of macrophages in our *in vitro* model and observed only a small decrease in bacterial growth rates that was independent of the virulence of the Ot strain indicating that the IFN-γ-mediated protection is not primarily due to increased bacterial killing by macrophages but rather the wider dysregulation in the immune response that was described in IFN-γ-deficient mice^46^. We showed that IFN-γ stimulation of mouse and human macrophages resulted in upregulation of numerous cytokines and chemokines, and these effects were augmented or supressed differently by different strains. Therefore, the exact effect of the downstream immune response directed by IFN-γ *in vivo* will differ substantially between strains, driving different disease outcomes.

What are the strain-specific characteristics that drive the observed differences in immune response and thus virulence? We discuss three possible factors here. First, Ot encodes a large arsenal of predicted effector proteins in both the Ank and TPR family. Whilst a core subset of seven Anks are found in all eight Ot genomes analysed to date, there are an additional 122 Anks present in only a subset of strains^49^. Anks are Type 1 secreted effector proteins that localize to various compartments of the host cell and interact with different partners. For example, Ank01 and Ank06 of strain Ikeda modulate NF-kB relocation to the nucleus^28^, whilst Ank13 of strain Ikeda is a nucleomodulin that modulates host gene transcription patterns^29^. The functions of most Ot Anks are unknown, but given the diversity in Ank repertoire between strains it is likely that these account for significant strain specific differences in their ability to manipulate host cell biology. For example, Ank13 was shown to modulate the expression of multiple host genes including those involved in the inflammatory response. Out of the strains involved in our study Ank13 is only present in Ikeda, Kato and UT76^24^. It is likely that there are other Ank proteins that also modulate different subsets of inflammatory genes and the exact combination of Anks with this activity present in a given strain will play a significant role in determining the host response to infection. The TPR proteins are less well understood then Anks, but the distribution of these effector proteins also differs significantly between strains. Together the scale and diversity of the Ank and TPR repertoire of Ot strains makes it highly likely that these play an important role in driving differential virulence. Second, despite a lack of obvious candidate virulence factors in the IR regions of the Ot genome, it is possible that strain specific differences in bacterial gene expression drive differences in host responses. Gene expression in bacteria is influenced by gene location within the genome, as differential supercoiling affects transcription and the gene’s proximity to the origin of replication impacts its copy number during DNA replication.The highly unusual inter-strain rearrangement of the Ot genome may therefore result in differences in gene expression despite conservation of promoter sequences, and these may play a role in differential interactions with host cells and therefore virulence. Third, whilst all Ot strains studied to date encode a core genome of 657 conserved genes in their IR regions, it is possible that different alleles of these genes drive different outcomes during infection. For example, active site mutations or changes in protein-protein interaction sites might lead to altered activity of a biological pathway that cannot be predicted from a simplistic analysis of the presence or absence of a particular gene. Together, these three factors offer some plausible explanations for the observed strain specific differences in virulence.

Our comparative analysis led to the identification of one clear phenotype that correlates with virulence: the subcellular position and ScaC expression pattern of the avirulent strain TA686. The altered subcellular position may reflect a straightforward difference in bacterial mechanism, for example altered intracellular trafficking driven by a difference in activity of ScaC. However, it may also reflect a more substantial difference in the host cell, driven by the presence of different effector proteins, such as differential localization or activity of host derived systems involved in anchoring Ot to the perinuclear region. Whilst the relationship between bacterial subcellular position and virulence is unclear, this is the first reported bacterial phenotype that correlates with a lack of virulence and therefore a detailed analysis of its relationship will be the subject of future study. One important limitation of this finding is that we only included one avirulent strain in our study due to a strong bias towards human isolates in available strain collections, and therefore we cannot comment on the generalizability of this finding. The availability of a larger collection of avirulent strains, for example including those isolated from mites, would enable larger scale comparisons to be made.

Our understanding of Ot cell biology has come from concerted efforts from multiple labs across the world. Historically, different research groups have worked on one or two Ot strains, often reflecting the geographic region where the research is based. One important conclusion from our study is that different Ot strains have different characteristics both at the level of *in vitro* cell biology and *in vivo* pathogenesis. Therefore, a greater emphasis on studying multiple strains would help the field to distinguish universal characteristics from strain specific features.

In summary, we have shown that different Ot strains exhibit differential virulence in a murine infection model and can be ranked into a hierarchy of virulence. We show that there is no single pattern of bacterial gene expression, *in vitro* growth rate, or inflammation that correlates with virulence, and we emphasize the importance of studying multiple different Ot strains. We show that the avirulent strain TA686 differs from other OT strains in its subcellular localization and ScaC expression, and hypothesize that this may be relevant in its differential virulence. We are hopeful that this study will help to drive forward the field of scrub typhus pathogenesis.

## Materials and Methods

### Cell Culture and Reagents

Bone marrow-derived macrophages (BMDMs) were isolated from 2-week-old male C57BL/6J mice and generously provided by Dr. Jelena Bezbradica Mirkovic. The murine monocytic cell line RAW264.7 was obtained from Dr. Samantha Bell. Mouse fibroblast cell line NCTC clone 929 (L-929) was purchased from ATCC (cat# CCL1).

BMDMs were cultured in DMEM-high glucose (Gibco, cat# 19650-62, Germany), supplemented with 10% fetal bovine serum (FBS; Gibco, cat# 16140071, USA), 1% Penicillin-Streptomycin-Glutamine (Gibco, cat# P4333, USA), 25 mM HEPES (Gibco, cat# 15630-056, USA), and 50 ng/ml Recombinant Mouse M-CSF (BioLegend, cat# 574806, USA). RAW264.7 and L929 cells were cultured in DMEM-high glucose, supplemented with 10% heat-inactivated FBS and 1% penicillin-streptomycin.

Human M1 macrophages (hMDM-GMCSF(-)) derived from a single donor were purchased from Promocell (cat# C-12914, Germany). These cells were cultured in M1-Macrophage Generation Medium XF (Promocell, cat# C-28055, Germany) according to the manufacturer’s instructions. The human monocytic cell line THP-1, provided by Dr. Jason Yang, was cultured in RPMI-1640 medium (Sigma-Aldrich, cat# R0883, USA) supplemented with 10% heat-inactivated FBS and 1% Penicillin-Streptomycin-Glutamine (Gibco, cat# 10378-016, USA). All cells were maintained at 37°C in a 5% CO2 atmosphere.

### Bacterial Strains and Propagation

Seven strains of *Orientia tsutsugamushi* were propagated in L929 cells following a previously described protocol^50^. Briefly, L929 cells were seeded in T75 flasks at a density of 3 × 10^6^ cells per flask and incubated overnight before infection. A frozen bacterial stock was thawed and added directly to the flasks. On day 6 post-infection, the medium was removed, and the infected cells were washed once with Phosphate Buffered Saline (PBS; Gibco, cat# 10010023, Germany). The cells were then scraped from the flask, resuspended in PBS, and lysed using a bead mill homogenizer (Fisher Scientific, Finland) with glass beads at a power setting of 5 for 1 minute. Cell debris and glass beads were removed by centrifugation at 800 × g for 3 minutes. The supernatant containing bacteria was transferred to new tubes and subjected to high-speed centrifugation at 14,000 × g for 10 minutes to pellet the bacteria. The bacterial pellet was resuspended in Sucrose-Phosphate-Glutamate Buffer (SPG) and stored at -80°C until use.

### Mouse Experiments

**Experiment 1.** This research was conducted in strict accordance with protocols approved by the Armed Forces Research Institute of Medical Sciences (AFRIMS) Animal Care and Use Committee, following Thai laws, the Animal Welfare Act, and all applicable U.S. Department of Agriculture, Office of Laboratory Animal Welfare, and U.S. Department of Defense guidelines. The approved protocol number was PN23-08. The animal research adhered to the Guide for the Care and Use of Laboratory Animals (8th Edition, NRC publication). AFRIMS is an AAALAC International-accredited facility located in Bangkok, Thailand.

Female C57BL/6NJcl mice (lot numbers 9-18 and 9-22) were obtained from Nomura Siam International Co., Ltd. (Bangkok, Thailand) and used at 6–9 weeks of age. The mice were housed under specific pathogen-free (SPF) conditions in an animal biosafety level 2 (ABSL2) facility at AFRIMS. Mice were co-housed (4 mice per case, 2 cases per group) in standard polycarbonate microisolator cages with filter tops and natural ventilation. Each cage was equipped with stainless steel feeding hoppers and water bottles, with temperature maintained at 21°C ± 1 and relative humidity between 30–70%.

Nine groups of mice (n=7 per group) were intravenously injected via the tail vein (using insulin syringes, 30G, ½”, BD Ultra-Fine II, Becton Dickinson) with 1 × 10⁷ genomic DNA copies of different *Orientia tsutsugamushi* strains (TA686, TA763, Gilliam, UT176, Karp, UT76, Ikeda, and Kato), as well as a mock-infection control. To account for variations in bacterial viability, in vitro microscopy and qPCR experiments were conducted on all bacterial stocks used to confirm that the number of viable organisms was comparable between strains. Mice were weighed daily, and clinical signs of morbidity were monitored and scored over the 8-day experimental period. Mice displaying signs of severe morbidity or >15% weight loss from baseline were humanely euthanized. At the end of the study, blood and tissue samples (brain, heart, lungs, kidneys, spleen, and liver) were collected for bacterial quantification and histopathology. Mice were euthanized using CO_2_ inhalation at a gas flow rate of 2 L/min at 15 psi CO_2_, maintained for at least 5 minutes after cessation of breathing. Death was confirmed by physical examination (absence of a heartbeat), followed by an adjunctive physical method such as cervical dislocation or exsanguination.

**Experiment 2.** This research was conducted at the Rutgers Public Health Research Institute BSL3 vivarium facility, in strict accordance with protocols approved by the Institutional Animal Care and Use Committee (IACUC) at Rutgers University. The approved protocol number was PROTO202000012.

Female C57BL/6NJ mice were obtained from The Jackson Laboratory (ME, USA) and used at 6– 9 weeks of age. The mice were housed under specific pathogen-free (SPF) conditions in an animal biosafety level 3 (ABSL3) facility.

Eight groups of mice (n=9 per group) were intravenously injected via the saphenous vein with 1.25 × 10^6^ genomic DNA copies of different *Orientia tsutsugamushi* strains (TA686, TA763, Gilliam, Karp, UT76, Ikeda, and Kato), as well as a mock-infection control. Mice were weighed daily, and clinical signs of morbidity were monitored and scored over the 12-day experimental period. Mice displaying signs of severe morbidity or >15% weight loss from baseline were humanely euthanized. At the end of the study, blood and tissue samples (brain, heart, lungs, kidneys, spleen, and liver) were collected for bacterial quantification and histopathology. Mice were euthanized via cervical dislocation and death was confirmed by prolonged observation and absence of heart beat.

### Histopathology Analysis

Spleen, lung, liver, kidney, heart and brain tissue were sectioned and prepared using hematoxylin and eosin stain. Histopathologic lesions were scored from 0-5, with 0 indicating normal tissue and 5 indicating severe inflammation. Scoring criteria: 0: normal, 1: minimal (possible background lesion), 2: mild, 3: moderate, 4: marked, 5: severe.

### Luminex Analysis

Mouse immune responses were assessed using the MILLIPLEX® multiplex assay (MCYTOMAG-70K, Merck Millipore) to measure nine analytes (GM-CSF, IFN-γ, IL-1α, IL-1β, IL-6, IL-10, IL-12(p70), IL-17, MCP-1), following the manufacturer’s instructions. Cytokine quantification was performed on a MAGPIX (Luminex) analyzer, and data were collected using Belysa® immunoassay software 1.2.2.

### *In Vitro* Growth Rate Analysis

Cells were seeded overnight at 1 or 3 × 10⁴ cells/well in 96- or 48-well plates, respectively. The following day, the medium was removed, and bacteria were added to the cells at a multiplicity of infection (MOI) of 50, followed by incubation for 3 hours. After incubation, bacteria were removed, and the infected cells were washed three times with plain medium. The cells were further cultured in growth medium under the same conditions. At specific time points, the medium was removed, and the infected cells were washed three times with PBS. DNA was extracted by adding alkaline lysis buffer (25 mM NaOH, 0.2 mM EDTA) directly into the wells, followed by boiling the plates at 95°C for 30 minutes to inactivate the bacteria. The plates were stored at 4°C until further testing.

Bacterial concentration was quantified by qPCR. The primers and TaqMan probe used for the 47 kDa target gene were as follows: 47 kDa FW (5’-TCCAGAATTAAATGAGAATTTAGGAC-3’), 47 kDa RV (5’-TTAGTAATTACATCTCCAGGAGCAA-3’), and 47 kDa probe (5’-FAM-TTCCACATTGTGCTGCAGATCCTTC-TAMRA-3’). The qPCR mixture consisted of 1X qPCR Probe Mix LO-ROX (PCR Biosystems, cat# PB20.21, UK), 0.1 µM of each forward and reverse primer, 0.2 µM probe, sterile distilled water, and 1 µL of extracted DNA. Real-time PCR was performed on a CFX Duet Thermal Cycler (Bio-Rad, USA) using the following conditions: initial denaturation at 95°C for 2 minutes, followed by 45 cycles of 95°C for 15 seconds and 60°C for 30 seconds, with fluorescence acquisition during the annealing/extension phase. DNA copy numbers were calculated by comparison with a standard curve.

### *In Vitro* Cytokine and Chemokine Expression Analysis

BMDM and human primary macrophages were seeded in 96-well plates at 1 × 10⁴ cells/well and cultured for 24 hours. Cells were either left unstimulated or stimulated for 18 hours with 400 U/ml of recombinant mouse (R&D Systems, cat# 485-MI-100, USA) or human IFN-γ (R&D Systems, cat# 285-IF-100, USA) for BMDM and human primary macrophages, respectively. Bacteria were then added in triplicate after 3 hours, and the infected cells were washed three times with plain medium. Infected cells were further cultured in the presence or absence of IFN-γ for 4 days. The cells were washed three times with PBS and lysed with alkaline lysis buffer. Bacterial concentration was quantified via real-time PCR, as previously described.

Host gene expression was determined by extracting total RNA from BMDM and human primary macrophages using the CellAmp Direct RNA Prep Kit (Takara, cat# 3732, Japan), following the manufacturer’s instructions. RNA was transcribed and amplified using the One Step TB Green® PrimeScript™ RT-PCR Kit (Takara, cat# RR066A, Japan). Relative quantitation of mRNA expression was calculated using the 2^(-ΔΔCt) method. Primers are listed in Supplementary Tables 1 and 2.

### Intracellular Infection Cycle and ScaC Expression Analysis

L929 cells were seeded and cultured in µ-Slide 8 Well plates (ibidi, cat# 80826, Germany) at 5 × 10³ cells/well for 24 hours. Bacteria were added at an MOI of 50 and incubated for 3 hours. After washing, cells were cultured for 4 and 7 days. On day 4, the number of cells with *Orientia tsutsugamushi* at different subcellular locations was counted, and at day 7, the expression of ScaC was analyzed using immunofluorescence staining. Cells were observed under a confocal microscope, and the data was manually analyzed.

### Statistical Analysis

All statistical analyses were performed using GraphPad Prism version 10.3.1.

## Acknowledgements

JS was supported by a Wellcome Trust Senior Research Fellowship (224277/Z/21/Z) and an NIH R21 award (R21AI144385).

Disclaimer: Material has been reviewed by the Walter Reed Army Institute of Research. There is no objection to its publication. The opinions or assertions contained herein are the private views of the author, and are not to be construed as official, or as reflecting true views of the Department of the Army or the Department of Defense. Research was conducted under an IACUC-approved animal use protocol in an AAALAC International - accredited facility with a Public Health Services Animal Welfare Assurance and in compliance with the Animal Welfare Act and other federal statutes and regulations relating to laboratory animals

---

**Supplementary Figure 1.** A. Results from high dose experiment in which C57Bl/6J were infected with 1x10^7^ bacteria per animal and monitored for 8 days. Kaplan-Meier survival curves of mice infected with different bacterial strains showing that only strain Kato causes lethal infection in this animal model. Statistical significance was determined using log-rank Mantel-Cox analysis.

## Notes

### Competing Interest Statement

The authors have declared no competing interest.

